# Molecular basis for the ATPase-powered substrate translocation by the Lon AAA+ protease

**DOI:** 10.1101/2020.04.29.068361

**Authors:** Shanshan Li, Kan-Yen Hsieh, Shih-Chieh Su, Grigore D. Pintilie, Kaiming Zhang, Chung-I Chang

## Abstract

The Lon AAA+ (adenosine triphosphatases associated with diverse cellular activities) protease (LonA) converts ATP-fuelled conformational changes into sufficient mechanical force to drive translocation of the substrate into a hexameric proteolytic chamber. To understand the structural basis for the substrate translocation process, we have determined the cryo-electron microscopy (cryo-EM) structure of *Meiothermus taiwanensis* LonA (MtaLonA) at 3.6 Å resolution in a substrate-engaged state. Substrate interactions are mediated by the dual pore-loops of the ATPase domains, organized in spiral staircase arrangement from four consecutive protomers in different ATP-binding and hydrolysis states; a closed AAA+ ring is nevertheless maintained by two disengaged ADP-bound protomers transiting between the lowest and highest position. The structure reveals a processive rotary translocation mechanism mediated by LonA-specific nucleotide-dependent allosteric coordination among the ATPase domains, which is induced by substrate binding.

## Introduction

The Lon AAA+ protease (LonA) is an ATP-dependent protease conserved in prokaryotes and eukaryotic organelles. LonA assembles as a homohexamer with each protomer containing an N-terminal domain, a middle ATPase domain, and a C-terminal protease domain (1). In addition to LonA, other Lon-like proteases (LonB and LonC) with distinct ATPase domains have been characterized in thermophilic archaeal and bacterial species (2, 3). LonA plays a major role in cellular protein homeostasis by degrading damaged or misfolded abnormal proteins, which prevents these unwanted protein species from forming toxic aggregates. LonA is also involved in the regulation of many biological processes by degrading specific regulatory proteins (4). Sharing the fused AAA+ and protease domain organization of Lon, the membrane-anchored AAA+ proteases FtsH and related mitochondrial intermembrane-space (i) and matrix (m) AAA (i/m-AAA) proteases are involved in quality control of membrane proteins; however, the ATPase domains belong to different clades of the superfamily and the protease domains are non-homologous (5, 6).

Previous results have revealed many functional and structural features of LonA. The protease activity of LonA is dependent on the presence of Mg^2+^, which binds to the protease domain and induces the formation of open active-site structure (7). The Lon protease domain with an open active-site conformation adopts a 6-fold symmetric hexameric ring (7–10). By contrast, LonA with an inactive proteolytic active-site conformation exhibits an open-spiral hexameric structure (11, 12). Therefore, the protease domain plays an important role in the closed-ring assembly of LonA. While ATP is required for LonA to degrade protein substrates efficiently, ADP is known to inhibit the protease activity of Lon. The crystal structure of the ADP-bound LonA hexamer has revealed how ADP may inhibit LonA activity by inducing the formation of a closed degradation chamber (13). A substrate-translocation model was posited based on the ADP-bound structure by assuming a structural transition between the nucleotide-free and ADP-bound states from two pairs of three non-neighboring protomers (13). However, the simple one-step transition model is not supported by recent cryo-EM structures of substrate-bound AAA+ proteins with the ATPase domains organized in spiral staircase arrangement around a centrally positioned substrate polypeptide chain (14–22). These structures support a processive rotary mechanism, which was also suggested by recent cryo-EM structures of human and *Yersinia pestis* Lon proteases (23, 24). However, in these works the bacterial Lon used for reconstruction was a Walker-B mutant; the structures of human and bacterial Lon show different engagement with the bound substrate from the six protomers, with different sets of nucleotide-binding states (23, 24).

Here, we report cryo-EM structures of *Meiothermus taiwanensis* LonA (MtaLonA) in a substrate-engaged state, determined at 3.6 Å resolution. Structural analysis suggests that both the substrate and ATP play an important role to induce a spiral staircase arrangement of the ATPase domains. Moreover, our structure of wild-type bacterial LonA reveals how binding of ATP and hydrolysis of ATP to ADP induce distinct nucleotide-induced conformational states in the six ATPase domains to enable a processive rotary translocation mechanism by LonA-specific allosteric coordination.

## Results

### Specimen preparation of LonA with a protein substrate

To capture MtaLonA in the substrate-engaged state, we assessed the degradation of Ig2 (25 kDa), a model substrate used previously (7, 25), by wild-type MtaLonA using the slowly hydrolyzable ATP analog, ATP-γ-S. At the optimal reaction temperature (55°C), MtaLonA is able to degrade Ig2 with ATP-γ-S, albeit much slower compared to ATP (**Figure S1**). The result suggests that Ig2 is productively translocated by MtaLonA utilizing ATP-γ-S. Therefore, for cryo-EM imaging, we purified MtaLonA in complex with Ig2, ATP-γ-S, and a peptidyl boronate, MMH8709 (13), which inhibits Lon protease activity by covalently binding to the catalytic serine in the protease domain, in order to trap MtaLonA in the process of translocating Ig2 without undergoing subsequent substrate proteolysis (7, 10)(See Methods for details).

### Cryo-EM structure of LonA engaged in translocating substrate

The structure of the MtaLonA-Ig2-MMH8709-ATP-γ-S complex was determined to 3.6-Å resolution by cryo-EM (**Figures 1A, S1-2, and Table S1**). The six protease domains associate into a closed-ring structure with C6 symmetry (**Figure 1B**). The six ATPase domains also form a closed ring (**Figure 1B**). However, those from four consecutive protomers P1-P4 (designated in the PDB coordinate file as chains A-D) organize into a spiral staircase arrangement, with protomers P1 and P4 occupying the highest and the lowest positions, respectively (**Figure 1C**). These four protomers spiral around an extended 11-residue polypeptide chain, likely representing a segment of unfolded substrate Ig2, of which the backbone is well resolved in the map. By contrast, the two relatively mobile promoters P5 and P6 (corresponding to chains E and F, respectively, in the PDB file) make no substrate contacts (**Figure 1D**). These non-engaged protomers are defined as the “seam” protomers as they detach from the staircase and make a loose association with each other (**Figures 1A–C**). Although the full-length construct of wild-type MtaLonA includes the N-terminal regions (residues 1-243), which is critical for substrate interaction and is required to degrade misfolded or native protein substrates (25), no density was found for these regions in our structures, likely due to their flexible nature.

**Figure 1.**
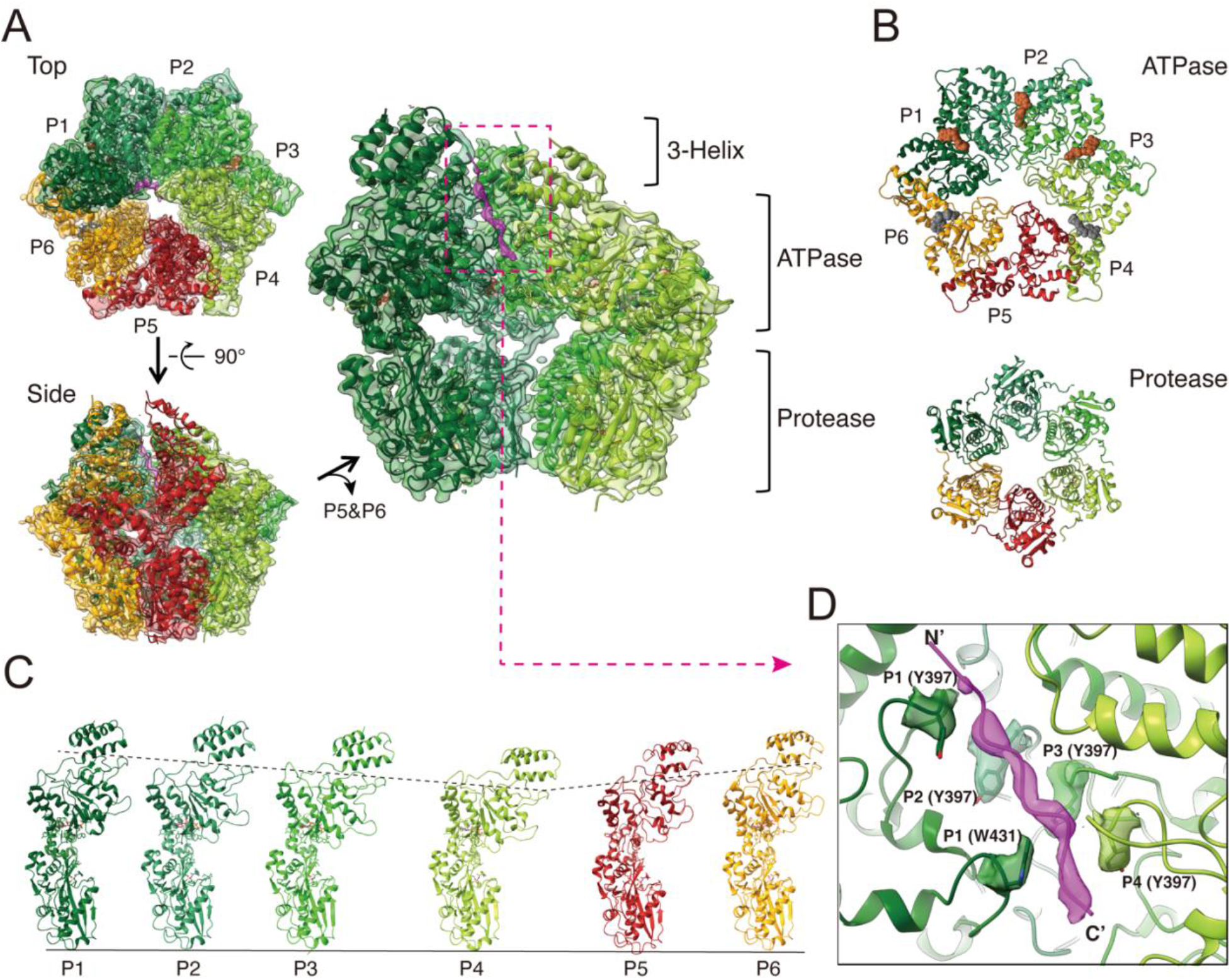
Overall structure of the substrate-bound MtaLonA complex. (A) The ribbon model of Ig2 (magenta) bound MtaLonA fitting into the 3.6-Å cryo-EM map (semi-transparent) is presented in the top and side views. The protomers in four substrate-engaged states (P1–P4) are colored in different shades of green; the protomers in two disengaged states (P5 and P6) are colored in brick red and orange, respectively. Protomers P5 and P6 of MtaLonA are removed in the close-up view (right) to show the density of Ig2 (dashed box) inside. (B) Top views show the rings of the ATPase domains (top) and the protease domains (bottom). ATP-γ-S and ADP are represented by brown and grey spheres, respectively. (C) Six conformational states (P1-6) of the protomers, arranged in clockwise order in the structure, are shown from left to right; the protomers are aligned using the protease domain as the reference. (D) Pore-loop 1 (PL1) and 2 (PL2) residues from the protomers P1-P4 interacting with Ig2 (magenta density) are shown with surrounding density. N’ and C’ denote the N- and C-terminal ends, respectively.

The sequence of Ig2 and the polarity of the bound polypeptide chain could not be determined from the density map. A strand of 11 alanine residues was modeled, with the C-terminus facing inside the chamber because Lon is known to recognize exposed C-terminal degron from model substrates and kinetics study indicates that the order of substrate scissile-site delivery occurs from the C-to N-terminal direction (26, 27). Contacts to the substrate polypeptide chain are made by protomers P1-P4, mediated by residue Tyr397 from pore-loop 1 and residue Trp431 from pore-loop 2; both have been shown to be required for degrading Ig2 (13). The four pore-loop 1 Tyr397 residues contact Ig2 in a right-handed spiral arrangement, with amino acid residues i, i+2, i+4, and i+6 of the substrate (**Figure 1D, Video S1**). Contrary to the well-resolved pore loops of substrate-engaged protomers P1-P4, those of the seam protomers P5 and P6 do not make contact with the substrate and are less well-resolved in the map (**Figures 1A and 1D**).

### ATPase sites around the substrate-engaged AAA+ ring

To understand how the staircase arrangement of the pore-loops 1 and 2 engaging the substrate correlates with nucleotide binding and hydrolysis states in the protomers, we analyzed the cryo-EM map of the ATPase site located in between the large AAA-α/β and the small AAA-α domains. The resolution of the map is sufficient to identify the bound nucleotides in the ATPase sites in protomers P1-P4 (**Figures S3, S4, and Video S1**). In protomer P1, the ATPase pocket is well ordered, with a bound ATP-γ-S, whose γ-phosphate interacts with the arginine finger (Arg finger) Arg484 from the clockwise neighboring protomer P2; the α- and β-phosphates are contacted by the sensor II, Arg541, in the cis protomer (**Figure 2A**). Similarly, in protomers P2 and P3, ATP-γ-S is found in the ATPase pocket with well-resolved density, with the γ-phosphate also contacted by the Arg finger. However, the sensor II Arg541 engages the β- and γ-phosphates of the bound ATP-γ-S in protomer P2 but solely with the γ-phosphate in protomer P3 (**Figures 2B and 2C**). As a result, the Arg finger and sensor II residues appear to present the scissile Pγ-Oβ bond of ATP-γ-S in the most suitable conformation for hydrolysis only in protomer P3. In contrast, the ATPase pocket of protomer P4, which occupies the lowest step of the substrate-bound staircase, is ADP-bound and more open than those of protomers P1-P3; the Arg finger from the protomer P5 is 15 Å away from the β-phosphate and the adjacent sensor II residue contacting α-phosphate is flexible (**Figure 2D**).

**Figure 2.**
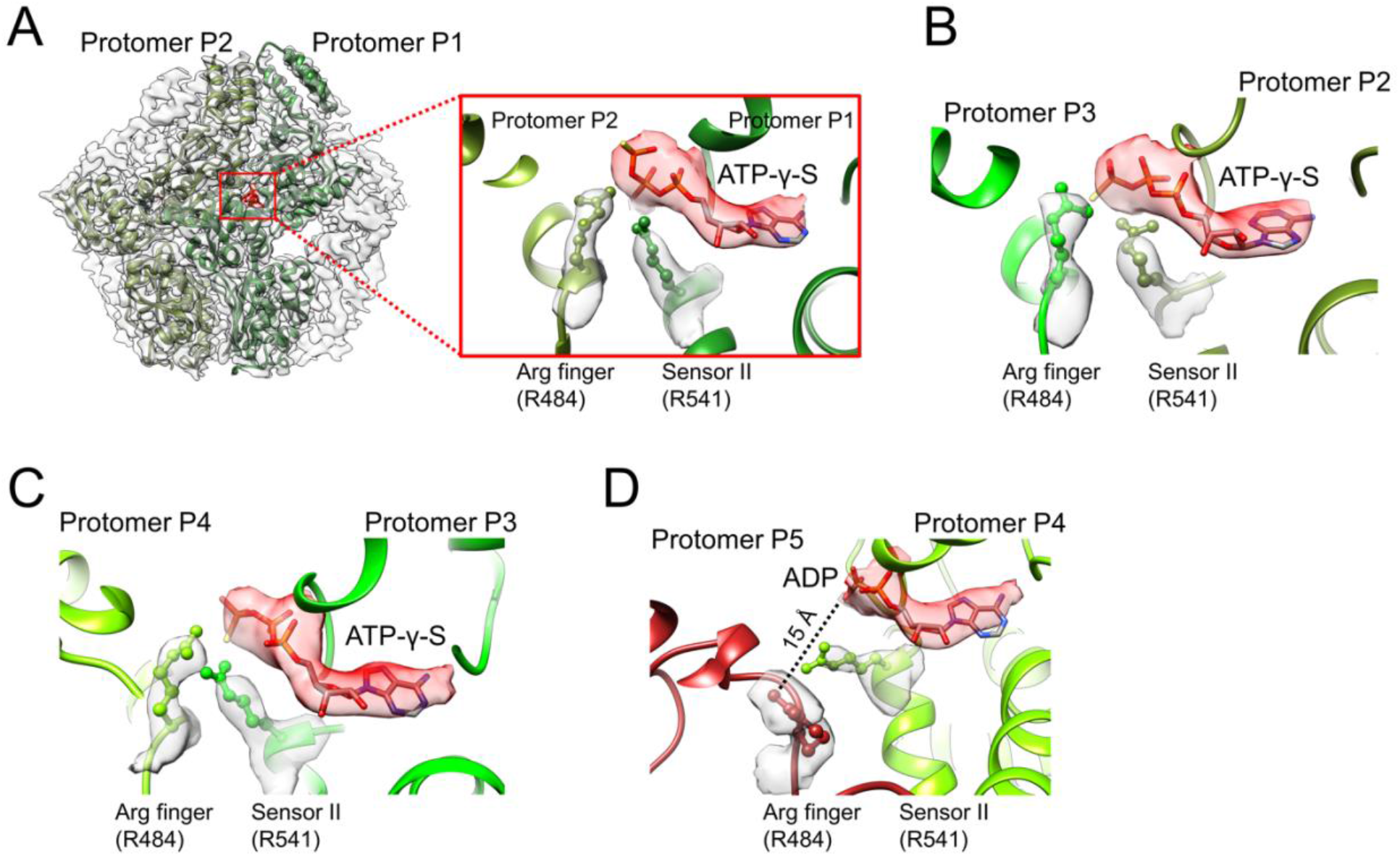
ATPase sites surrounding the substrate-engaged AAA+ ring. (A) The arginine finger and sensor II residues interacting with the ATP-γ-S bound to protomer P1. (B) The arginine finger and sensor II residues interacting with the ATP-γ-S bound to protomer P2. (C) The arginine finger and sensor II residues interacting with the ATP-γ-S bound to protomer P3. (D) The arginine finger and sensor II residues do not interact with ADP bound to protomer P4. The bound nucleotide is shown in red density with the fitted model.

Interestingly, the AAA-α/β domain of the disengaged seam protomer P5 makes the least inter-protomer interaction and is highly mobile as indicated in the resolution map (**Figure S2E**) and Q-scores (**Figure S4**), which measures the local resolution (28) and structure resolvability (29), respectively. The ATPase site has only broken density for the bound nucleotide and the Walker-A/B motifs (**Figure S3**); the Arg finger from protomer P6 and the sensor II are also unresolved. As mentioned above, the Arg finger R484 from protomer P5 is far from the bound ADP in protomer P4; therefore, the protomer P5 ATPase site is likely bound to ADP also, but not ATP-γ-S.

The seam protomer P6 also has a mobile AAA-α/β domain. The resolution of the map at the ATPase pocket is nevertheless sufficient to reveal a bound ADP (**Figure S3**), probably due to its ordered neighboring protomer P1. The side chains of the Arg finger from protomer P1 and the sensor II residues, though located nearby, are not well resolved.

### Substrate-induced allosteric coordination

In the substrate-engaged state, the spiral staircase arrangement of the AAA-α/β domains in the closed AAA+ ring, which is covalently fused to the closed protease ring, is made possible by several previously found LonA-specific structural features (13). They include nucleotide-dependent rigid-body rotation of the AAA-α/β domain, which presents the dual pore-loops, to adopt different positions with respect to the AAA-α domain. In addition, the flexible protease-domain (PD) linker bridging the AAA-α and the protease domains accommodate further rotational movement of the AAA-α domain (**Figure 3**).

**Figure 3.**
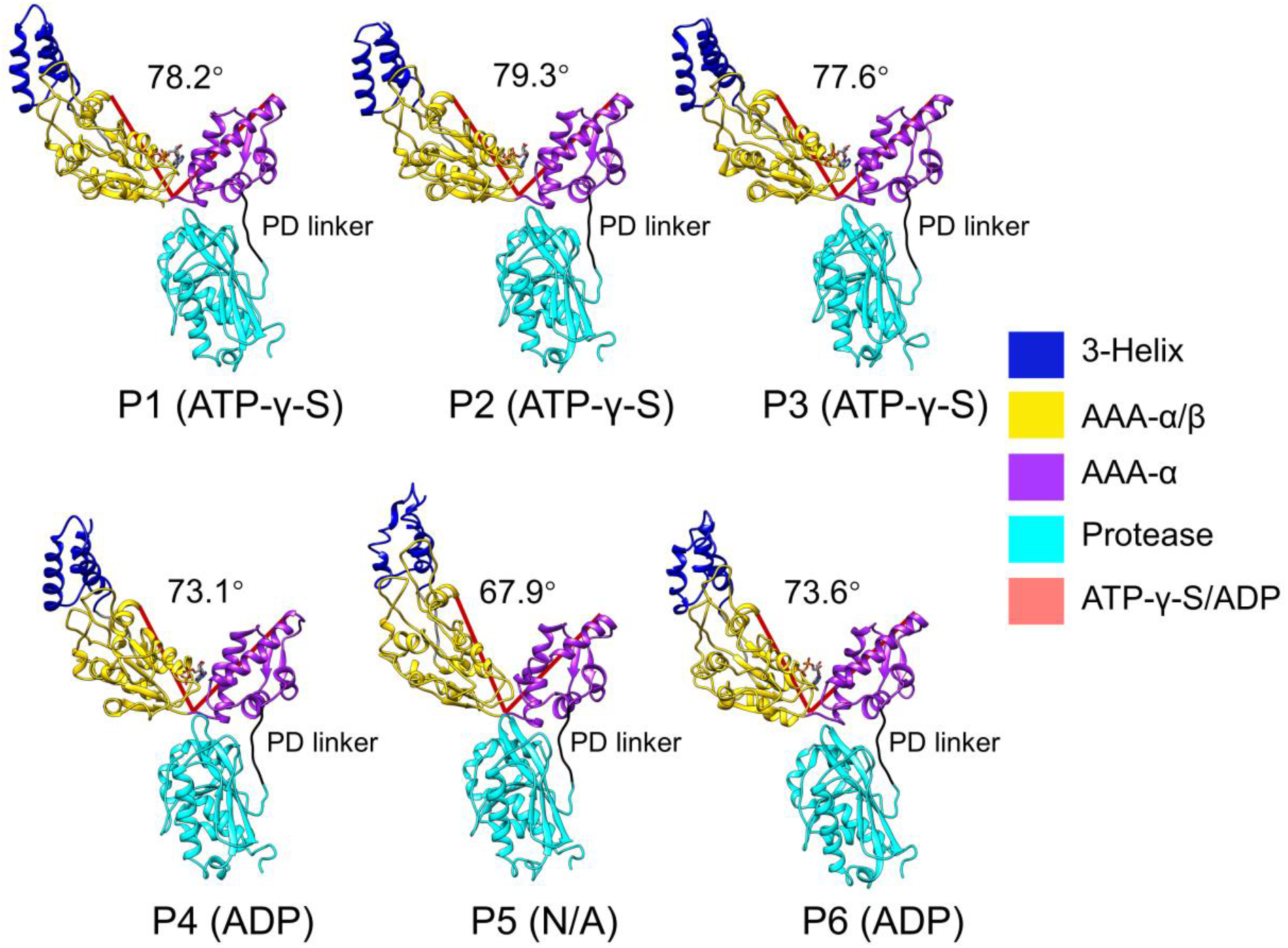
Nucleotide-dependent rigid-body rotation of the AAA-α/β domain. The protomers were aligned based on the AAA-α domain. The dihedral angles between the AAA-α/β and AAA-α domains are shown by using the Ca atom of Leu559, Gly492, and Gln411 as reference points for measurement. The protease-domain (PD) linker (a.a. 584-589) is shown in black.

Notably, the cis Arg finger Arg484 in the ATP-γ-S-bound protomers are positioned within the hydrogen-bonding distance near the bound nucleotide in the ATPase pocket of the counterclockwise neighboring protomer, making contact only with bound ATP-γ-S but not ADP (**Figure 2**). Interestingly, Arg484 is contacted by Asp444 and Pro445 in the N-terminal base-loop of the pre-sensor-1 β-hairpin (PS1βH), a motif conserved in LonA, HslU, and Clp AAA+ proteases (30, 31). In the substrate-engaged, spirally arranged protomers P1-P3, the base-loop residues Asp444 and Glu446 form a trans-ATP-interacting group together with Arg484 to bind ATP-γ-S (**Figure S5**). Taken together, based on the structural results it may be assumed that interaction with substrate polypeptide chain appears to induce the formation of a spiral staircase arrangement of ATP-bound protomers, enabling the interaction of the PS1βH base-loop and the Arg-finger with the ribose and the phosphate groups of ATP, respectively, and to facilitate ATP hydrolysis in the ATPase site. Our structure may thus offer a straightforward explanation of how the presence of protein or peptide substrates stimulates the ATPase activity of LonA, a hallmark feature also shared by other AAA+ proteases (32–37).

## Discussion

In this work, we have determined the structure of MtaLonA in the act of translocating the model substrate Ig2, which is partially folded protein (25). The substrate-bound structure forms an asymmetric close ring. Interestingly, recent cryo-EM structures of substrate-free Lon show an open spiral structure, where the side opening potentially allows access of the pore-loops to substrates from outside (11, 23, 24). Nevertheless, the substrate Ig2 is more likely to be translocated from the top of the AAA+ ring, where the substrate-binding N-terminal domain, essential for recognition and proteolysis of Ig2 (25), is located.

The main goal of the present work is to address the molecular basis for substrate translocation by LonA. Our structure of Ig2-bound wild-type MtaLonA suggests a LonA-specific processive rotary mechanism to drive translocation of substrate polypeptide chains (**Figure 4**). As shown in the schematic diagrams, during the translocation process a polypeptide chain is gripped by four protomers, each of these protomers traverses in clockwise direction through successive cycles of six conformational states, of which four are in substrate-engaged modes and two are in non-engaged detached modes. Of the four engaged modes, two are coordinated by binding of ATP, one presumably triggers ATP hydrolysis (indicated by the lightning symbol), and one is bound to ADP. Protomers of LonA in the engaged states communicate with one another by the Arg finger and the PS1βH base-loop. In the two detached modes, the protomers adopt mobile conformations permitting exchange of ADP by ATP, after which they are switched sequentially back into the substrate-engaged mode. In the next cycle, ATP hydrolysis proceeds counterclockwise to the next ATPase site.

**Figure 4.**
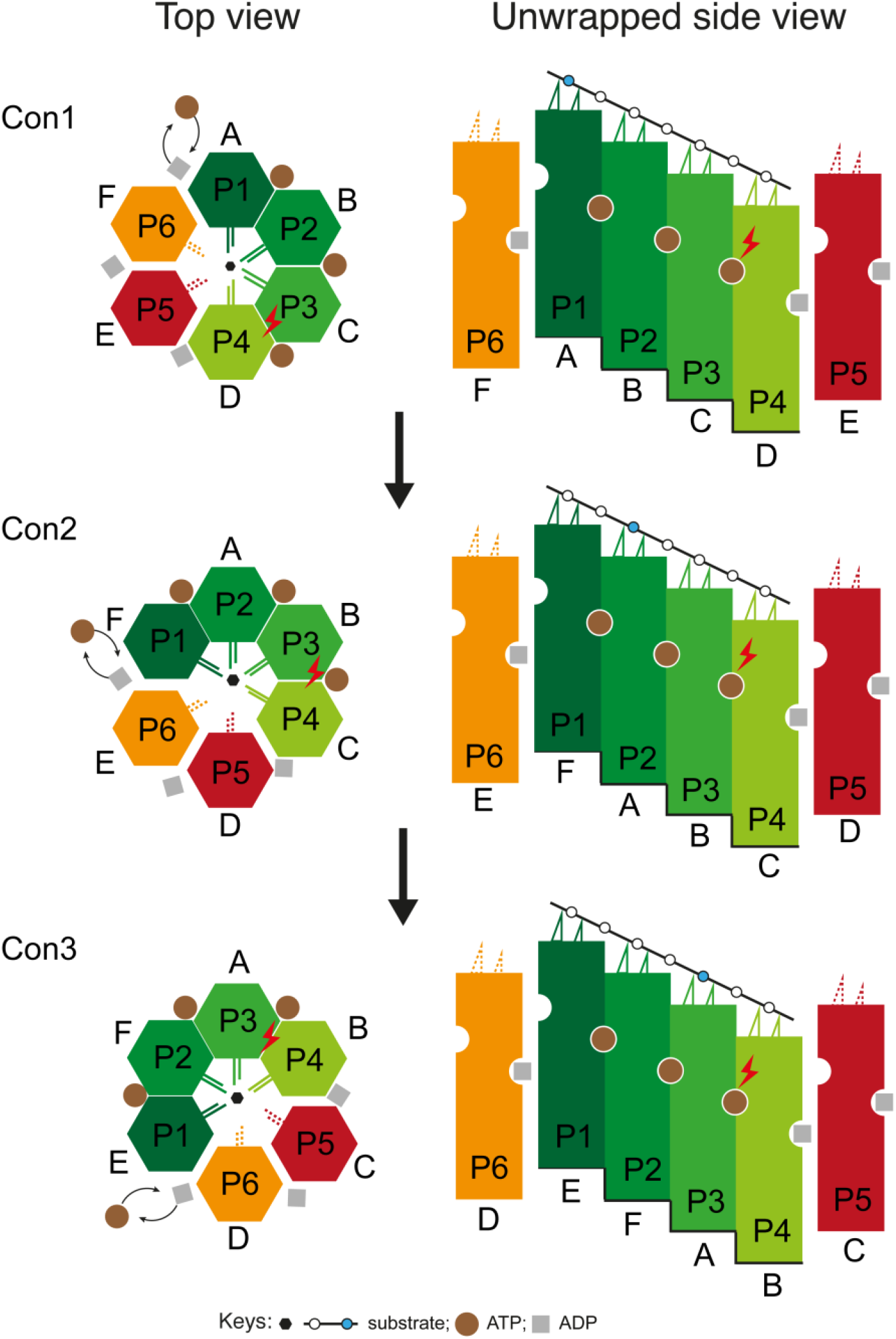
Structure-based model proposed for the processive substrate translocation by MtaLonA. Sequential ATP hydrolysis steps are shown in the top-view (left) and unwrapped side-view (right) diagrams. In the top view diagram, the colored hexagons represent the six conformational states (P1~P6), which in Con1 are adopted by specific protomer chains (A~F) as shown in the cryo-EM structure. Protomers in four substrate-engaging states (P1~4) are colored in different shades of green. Protomers in the non-engaging states P5 and P6 are shown as detached hexagons colored in red and orange, respectively. The protomer chains (A~F) are fixed in position to show that substrate engagement, ATP hydrolysis, and nucleotide exchange proceed in the ATPase ring counterclockwise during the transition from Con1 to Con3. The side-view diagram shows the staircase arrangement formed by the conformational states (P1~P4) of the protomers, represented by vertical rectangles, with the notches on the left and right sides of each rectangle representing the trans-ATP-interacting and the ATPase sites, respectively (see text). During the transition from Con1 to Con3, the staircase arrangement is adopted by different sets of protomer chains (A~F), of which two adopt non-engated states shown as detached rectangles. The two pore-loops of each protomer are depicted by the double lines in the top view and the triangles in the side view. Dashed double lines denote flexible pore loops. Putative ATP hydrolysis sites are depicted by lightning symbols. Polypeptide chain in the side-view diagram is denoted by a straight line with small circles marking each of the amino acid residues. Translocation of the substrate is depicted by the movement of an arbitrarily chosen residue colored in blue, which binds to chain A going through successive conformational states.

The mechanism featuring “hand-over-hand” substrate translocation coupled to counterclockwise sequential hydrolysis of ATP is conserved in a variety of different AAA+ proteins (14–22). A recent cryo-EM study describing the structure of a substrate-bound Walker-B mutant of LonA from *Yersinia pestis* (YpeLonA) also suggested a similar mechanism (24). However, these substrate-bound structures show several notable differences. Despite containing four ATP-bound protomers in the staircase arrangement, YpeLonA engaged the substrate via only the highest three spiraling pore-loop 1 residues; in addition, the structure did not show direct interaction of the substrate with pore-loop 2. By contrast, MtaLonA binds to the substrate through the pore-loops from all four staircase protomers, with the lowest protomer of the staircase bound to ADP; such nucleotide arrangement in the substrate-engaging protomers has also been observed in the structure of a DNA-engaged state of the CMG helicase (38).

Comparison of the LonA structures with other similar AAA+ proteases also showed notable differences in the nucleotide-binding and substrate-engaging states (Figure S6) (16, 22, 39–41). Notably, results from these AAA+ proteases show that translocation may be driven by engaging substrate polypeptides with 3~5 protomers. Therefore, although operated by a common mechanism, different AAA+ proteases may respond to different substrates by employing different numbers of protomers for translocation; participating protomers traverse through different conformational states to coordinate ATP hydrolysis, which may generate different mechanical pulling forces. In this regard, future investigations on the structures of AAA+ proteases with a series of substrates in different folding states may yield further insights into the mechanism of substrate unfolding and translocation by these proteolytic machines.

## Experimental procedures

### Protein expression and purification

Cells were grown in LB medium at 37°C until the optical density reached 0.6 to 0.8. Isopropyl β-d-thiogalactopyranoside (IPTG) to a final concentration of 1 mM was then added to the culture and incubated for another 4 h at 25°C for full-length MtaLonA and 37°C for the substrate Ig2 (7, 13). Cells were harvested by centrifugation and suspended in lysis buffer containing 50 mM Tris-HCl (pH 8.0) and 500 mM NaCl. Cell lysate was first ruptured by a French press (Avestin) and centrifuge at 35,000 g. Supernatant was collected for 2.5 h binding with Ni-nitrilotriacetic acid resins (Qiagen) at 4°C. Protein was further washed with a stepwise imidazole gradient and eluted with 250 mM imidazole. After that, Ig2 was treated with TEV protease overnight to remove the 6xHis tag. All proteins were dialyzed against different buffer components to remove imidazole. MtaLonA was dialyzed against buffer containing 20 mM Tris-HCl (pH 8.0), 100 mM NaCl and 2 mM DTT, while Ig2 was dialyzed against 25 mM Tris-HCl (pH 8.0), 150 mM NaCl and 2 mM β-mercaptoethanol. The untagged Ig2 was further purified by Ni-NTA and Superdex 200 (GE Healthcare) chromatography.

After purification, MtaLonA was first incubated with 10 mM MgCl2 for 1 h and then incubated with 1 mM MMH8709 (13), 5 mM adenosine 5′-[γ-thio]-triphosphate (ATP-γ-S) and Ig2 (in 5-fold molar excess) overnight. The MtaLonA-Ig2-MMH8709-ATP-γ-S complex was then loaded onto Superdex 200 (GE Healthcare) column pre-equilibrated with 20 mM Tris-HCl (pH8.0), 100 mM NaCl, 10 mM MgCl2 and 2 mM DTT to remove unbound compounds. The protein complex was treated with additional 1 mM of ATP-γ-S before cryo-sample preparation.

### Cryo-EM data acquisition

The substrate-engaged MtaLonA sample was diluted at a final concentration of around 0.15 mg/mL. Three microliters of the sample was applied onto glow-discharged 200-mesh R1.2/1.3 Quantifoil grids. The grids were blotted for 4 s and rapidly cryocooled in liquid ethane using a Vitrobot Mark IV (Thermo Fisher Scientific) at 4°C and 100% humidity. They were screened using a Talos Arctica cryo-electron microscope (Thermo Fisher Scientific) operated at 200 kV. One good grid was then imaged in a Titan Krios cryo-electron microscope (Thermo Fisher Scientific) with GIF energy filter (Gatan) at a magnification of 215,000× (corresponding to a calibrated sampling of 0.65 Å per pixel). Micrographs were recorded by EPU software (Thermo Fisher Scientific) with a Gatan K2 Summit direct electron detector, where each image was composed of 30 individual frames with an exposure time of 6 s and an exposure rate of 8.5 electrons per second per Å^2^. A total of 1,800 movie stacks were collected.

### Single-particle image processing and 3D reconstruction

All micrographs were first imported into Relion (42) for image processing. The motion-correction was performed using MotionCor2 (43) and the contrast transfer function (CTF) was determined using CTFFIND4 (44). All particles were autopicked using the NeuralNet option (threshold 1 = 0; threshold 2 = −5) in EMAN2 (45), and further checked manually. Then, particle coordinates were imported to Relion, where the poor 2D class averages were removed by several rounds of 2D classification. A total of 102,419 particles were transferred to cryoSPARC (46) for ab-initio map generation. Then a good class with 58,276 particles was imported to Relion for 3D classification. After removing bad classes, the final 3D refinement was performed using 23,487 particles, and a 3.6 Å map was obtained.

### Model building

The crystal structure of a hexameric LonA protease (PDB ID: 4YPL) from our previous work (13) was rigidly fitted into the cryo-EM map of substrate-engaged MtaLonA. As the six protomers were conformationally different in the cryo-EM density, molecular dynamics flexible fitting (MDFF) (47) was used. The MDFF was completed in three runs, where each run included 10^4^ minimization steps and 10^5^ molecular dynamics steps. After no noticeable deformation, the MDFF was stopped. The resultant models were refined using phenix.real_space_refine (48). The type of the bound nucleotide, ADP or ATP-γ-S, was determined by LigandFit in Phenix, with an overall correlation coefficient of the ligand to the map over 0.7. Then phenix.real_space_refine and Coot (49) were applied for model optimization.

The final model was evaluated by MolProbity (50). Statistics of the map reconstruction and model building are summarized in Table S1. The final structure model was analyzed with the PDBsum structure bioinformatics software (51) to identify key residues that interact with bound nucleotides. All figures were prepared using PyMol (52) and Chimera (53).

### Substrate degradation assay

Ig2 (domains 5 and 6 of the gelation factor ABP-120 of *Dictyostelium discoideum (54)*) was used as the substrate in this assay. Then 4 mM of the substrate protein was incubated with 0.8 mM MtaLonA (hexamer) in the reaction buffer containing 50 mM Tri-HCl pH 8.0, 10 mM MgCl2, 1 mM DTT, 5 mM ATP or ATP-γ-S at 55°C. At different time points, reaction aliquots were stopped by adding 5X SDS-PAGE loading dye and heated at 95°C for 5 min. The samples were then loaded onto a SurePAGE gel (4-20% Bis-Tris)(Genscript). Substrate protein bands were detected by Coomassie Blue staining.

## Acknowledgments

We thank Kuei-Hsiang Pan and Lu-Chu Ke for their help on protein preparation. This work was supported by Academia Sinica and MOST grant 108-2320-B-001-011-MY3 (to C.I.C.) and the Start-up funding from University of Science and Technology of China (KY9100000032 and KJ2070000080 to K.M.Z.).

## Author contributions

K.Z. and C.I.C. conceived the study and designed experiments; S.L. and K.Z. solved the structures; S.L., K.Y.H., and K.Z. performed experiments; S.L., K.Y.H., G.D.P., K.Z., and C.I.C. analyzed data; and S.L., K.Y.H., K.Z., and C.I.C. wrote the manuscript.

## Data deposition and availability

The cryo-EM map of the substrate-engaged MtaLonA with its associated atomic model have been deposited in the Electron Microscopy Data Bank and the Protein Data Bank under accession code EMD-21870, and PDB ID code 6WQH. All data needed to evaluate the conclusions in the paper are present in the paper and the supporting Information.

## References

1. Rotanova, T. V., Botos, I., Melnikov, E. E., Rasulova, F., Gustchina, A., Maurizi, M. R., and Wlodawer, A. (2006) Slicing a protease: structural features of the ATP-dependent Lon proteases gleaned from investigations of isolated domains. Protein Sci. 15, 1815–1828

2. Liao, J.-H., Kuo, C.-I., Huang, Y.-Y., Lin, Y.-C., Lin, Y.-C., Yang, C.-Y., Wu, W.-L., Chang, W.-H., Liaw, Y.-C., Lin, L.-H., Chang, C.-I., and Wu, S.-H. (2012) A Lon-like protease with no ATP-powered unfolding activity. PLoS One. 7, e40226

3. Rotanova, T. V., Melnikov, E. E., Khalatova, A. G., Makhovskaya, O. V., Botos, I., Wlodawer, A., and Gustchina, A. (2004) Classification of ATP-dependent proteases Lon and comparison of the active sites of their proteolytic domains. Eur. J. Biochem. 271, 4865–4871

4. Gur, E. (2013) The Lon AAA+ protease. Subcell. Biochem. 66, 35–51

5. Puchades, C., Sandate, C. R., and Lander, G. C. (2020) The molecular principles governing the activity and functional diversity of AAA+ proteins. Nat. Rev. Mol. Cell Biol. 21, 43–58

6. Sauer, R. T., and Baker, T. A. (2011) AAA+ proteases: ATP-fueled machines of protein destruction. Annu. Rev. Biochem. 80, 587–612

7. Su, S.-C., Lin, C.-C., Tai, H.-C., Chang, M.-Y., Ho, M.-R., Babu, C. S., Liao, J.-H., Wu, S.-H., Chang, Y.-C., Lim, C., and Chang, C.-I. (2016) Structural Basis for the Magnesium-Dependent Activation and Hexamerization of the Lon AAA+ Protease. Structure. 24, 676–686

8. Botos, I., Melnikov, E. E., Cherry, S., Tropea, J. E., Khalatova, A. G., Rasulova, F., Dauter, Z., Maurizi, M. R., Rotanova, T. V., Wlodawer, A., and Gustchina, A. (2004) The Catalytic Domain ofEscherichia coliLon Protease Has a Unique Fold and a Ser-Lys Dyad in the Active Site. Journal of Biological Chemistry. 279, 8140–8148

9. Cha, S.-S., An, Y. J., Lee, C. R., Lee, H. S., Kim, Y.-G., Kim, S. J., Kwon, K. K., De Donatis, G. M., Lee, J.-H., Maurizi, M. R., and Kang, S. G. (2010) Crystal structure of Lon protease: molecular architecture of gated entry to a sequestered degradation chamber. EMBO J. 29, 3520–3530

10. Liao, J.-H., Ihara, K., Kuo, C.-I., Huang, K.-F., Wakatsuki, S., Wu, S.-H., and Chang, C.-I. (2013) Structures of an ATP-independent Lon-like protease and its complexes with covalent inhibitors. Acta Crystallogr. D Biol. Crystallogr. 69, 1395–1402

11. Botos, I., Lountos, G. T., Wu, W., Cherry, S., Ghirlando, R., Kudzhaev, A. M., Rotanova, T. V., de Val, N., Tropea, J. E., Gustchina, A., and Wlodawer, A. (2019) Cryo-EM structure of substrate-free E. coli Lon protease provides insights into the dynamics of Lon machinery. Current Research in Structural Biology. 1, 13–20

12. Duman, R. E., and Löwe, J. (2010) Crystal structures of Bacillus subtilis Lon protease. J. Mol. Biol. 401, 653–670

13. Lin, C.-C., Su, S.-C., Su, M.-Y., Liang, P.-H., Feng, C.-C., Wu, S.-H., and Chang, C.-I. (2016) Structural Insights into the Allosteric Operation of the Lon AAA+ Protease. Structure. 24, 667–675

14. Monroe, N., Han, H., Shen, P. S., Sundquist, W. I., and Hill, C. P. (2017) Structural basis of protein translocation by the Vps4-Vta1 AAA ATPase. Elife. 10.7554/eLife.24487

15. Yu, H., Lupoli, T. J., Kovach, A., Meng, X., Zhao, G., Nathan, C. F., and Li, H. (2018) ATP hydrolysis-coupled peptide translocation mechanism of Mycobacterium tuberculosis ClpB. Proceedings of the National Academy of Sciences. 115, E9560–E9569

16. de la Peña, A. H., Goodall, E. A., Gates, S. N., Lander, G. C., and Martin, A. (2018) Substrate-engaged 26 proteasome structures reveal mechanisms for ATP-hydrolysis-driven translocation. Science. 10.1126/science.aav0725

17. Ripstein, Z. A., Huang, R., Augustyniak, R., Kay, L. E., and Rubinstein, J. L. (2017) Structure of a AAA+ unfoldase in the process of unfolding substrate. Elife. 10.7554/eLife.25754

18. Ho, C.-M., Beck, J. R., Lai, M., Cui, Y., Goldberg, D. E., Egea, P. F., and Zhou, Z. H. (2018) Malaria parasite translocon structure and mechanism of effector export. Nature. 561, 70–75

19. Gates, S. N., Yokom, A. L., Lin, J., Jackrel, M. E., Rizo, A. N., Kendsersky, N. M., Buell, C. E., Sweeny, E. A., Mack, K. L., Chuang, E., Torrente, M. P., Su, M., Shorter, J., and Southworth, D. R. (2017) Ratchet-like polypeptide translocation mechanism of the AAA+ disaggregase Hsp104. Science. 357, 273–279

20. White, K. I., Zhao, M., Choi, U. B., Pfuetzner, R. A., and Brunger, A. T. (2018) Structural principles of SNARE complex recognition by the AAA+ protein NSF. Elife. 10.7554/eLife.38888

21. Han, H., Monroe, N., Sundquist, W. I., Shen, P. S., and Hill, C. P. (2017) The AAA ATPase Vps4 binds ESCRT-III substrates through a repeating array of dipeptide-binding pockets. Elife. 10.7554/eLife.31324

22. Ripstein, Z. A., Vahidi, S., Houry, W. A., Rubinstein, J. L., and Kay, L. E. (2020) A processive rotary mechanism couples substrate unfolding and proteolysis in the ClpXP degradation machinery. Elife. 10.7554/eLife.52158

23. Shin, M., Watson, E. R., Song, A. S., Mindrebo, J. T., Novick, S. J., Griffin, P. R., Wiseman, R. L., and Lander, G. C. (2021) Structures of the human LONP1 protease reveal regulatory steps involved in protease activation. Nat. Commun. 12, 3239

24. Shin, M., Puchades, C., Asmita, A., Puri, N., Adjei, E., Wiseman, R. L., Karzai, A. W., and Lander, G. C. (2020) Structural basis for distinct operational modes and protease activation in AAA+ protease Lon. Sci Adv. 6, eaba8404

25. Tzeng, S.-R., Tseng, Y.-C., Lin, C.-C., Hsu, C.-Y., Huang, S.-J., Kuo, Y.-T., and Chang, C.-I. (2021) Molecular insights into substrate recognition and discrimination by the N-terminal domain of Lon AAA+ protease. Elife. 10.7554/eLife.64056

26. Mikita, N., Cheng, I., Fishovitz, J., Huang, J., and Lee, I. (2013) Processive degradation of unstructured protein by Escherichia coli Lon occurs via the slow, sequential delivery of multiple scissile sites followed by rapid and synchronized peptide bond cleavage events. Biochemistry. 52, 5629–5644

27. Gur, E., and Sauer, R. T. (2008) Recognition of misfolded proteins by Lon, a AAA protease. Genes & Development. 22, 2267–2277

28. Kucukelbir, A., Sigworth, F. J., and Tagare, H. D. (2014) Quantifying the local resolution of cryo-EM density maps. Nat. Methods. 11, 63–65

29. Pintilie, G., Zhang, K., Su, Z., Li, S., Schmid, M. F., and Chiu, W. (2020) Measurement of atom resolvability in cryo-EM maps with Q-scores. Nat. Methods. 17, 328–334

30. Iyer, L. M., Leipe, D. D., Koonin, E. V., and Aravind, L. (2004) Evolutionary history and higher order classification of AAA+ ATPases. J. Struct. Biol. 146, 11–31

31. Erzberger, J. P., and Berger, J. M. (2006) Evolutionary relationships and structural mechanisms of AAA+ proteins. Annu. Rev. Biophys. Biomol. Struct. 35, 93–114

32. Cheng, I., Mikita, N., Fishovitz, J., Frase, H., Wintrode, P., and Lee, I. (2012) Identification of a region in the N-terminus of Escherichia coli Lon that affects ATPase, substrate translocation and proteolytic activity. J. Mol. Biol. 418, 208–225

33. Goldberg, A. L., Moerschell, R. P., Chung, C. H., and Maurizi, M. R. (1994) ATP-dependent protease La (lon) from Escherichia coli. Methods Enzymol. 244, 350–375

34. Seol, J. H., Yoo, S. J., Shin, D. H., Shim, Y. K., Kang, M. S., Goldberg, A. L., and Chung, C. H. (1997) The heat-shock protein HslVU from Escherichia coli is a protein-activated ATPase as well as an ATP-dependent proteinase. Eur. J. Biochem. 247, 1143–1150

35. Waxman, L., and Goldberg, A. L. (1986) Selectivity of intracellular proteolysis: protein substrates activate the ATP-dependent protease (La). Science. 232, 500–503

36. Yamada-Inagawa, T., Okuno, T., Karata, K., Yamanaka, K., and Ogura, T. (2003) Conserved pore residues in the AAA protease FtsH are important for proteolysis and its coupling to ATP hydrolysis. J. Biol. Chem. 278, 50182–50187

37. Zhang, X., and Wigley, D. B. (2008) The “glutamate switch” provides a link between ATPase activity and ligand binding in AAA+ proteins. Nat. Struct. Mol. Biol. 15, 1223–1227

38. Eickhoff, P., Kose, H. B., Martino, F., Petojevic, T., Abid Ali, F., Locke, J., Tamberg, N., Nans, A., Berger, J. M., Botchan, M. R., Yardimci, H., and Costa, A. (2019) Molecular Basis for ATP-Hydrolysis-Driven DNA Translocation by the CMG Helicase of the Eukaryotic Replisome. Cell Rep. 28, 2673–2688.e8

39. Dong, Y., Zhang, S., Wu, Z., Li, X., Wang, W. L., Zhu, Y., Stoilova-McPhie, S., Lu, Y., Finley, D., and Mao, Y. (2019) Cryo-EM structures and dynamics of substrate-engaged human 26S proteasome. Nature. 565, 49–55

40. Puchades, C., Rampello, A. J., Shin, M., Giuliano, C. J., Wiseman, R. L., Glynn, S. E., and Lander, G. C. (2017) Structure of the mitochondrial inner membrane AAA+ protease YME1 gives insight into substrate processing. Science. 10.1126/science.aao0464

41. Puchades, C., Ding, B., Song, A., Wiseman, R. L., Lander, G. C., and Glynn, S. E. (2019) Unique Structural Features of the Mitochondrial AAA+ Protease AFG3L2 Reveal the Molecular Basis for Activity in Health and Disease. Mol. Cell. 75, 1073–1085.e6

42. Scheres, S. H. W. (2012) RELION: implementation of a Bayesian approach to cryo-EM structure determination. J. Struct. Biol. 180, 519–530

43. Zheng, S. Q., Palovcak, E., Armache, J.-P., Verba, K. A., Cheng, Y., and Agard, D. A. (2017) MotionCor2: anisotropic correction of beam-induced motion for improved cryo-electron microscopy. Nat. Methods. 14, 331–332

44. Rohou, A., and Grigorieff, N. (2015) CTFFIND4: Fast and accurate defocus estimation from electron micrographs. J. Struct. Biol. 192, 216–221

45. Tang, G., Peng, L., Baldwin, P. R., Mann, D. S., Jiang, W., Rees, I., and Ludtke, S. J. (2007) EMAN2: an extensible image processing suite for electron microscopy. J. Struct. Biol. 157, 38–46

46. Punjani, A., Rubinstein, J. L., Fleet, D. J., and Brubaker, M. A. (2017) cryoSPARC: algorithms for rapid unsupervised cryo-EM structure determination. Nat. Methods. 14, 290–296

47. Trabuco, L. G., Villa, E., Mitra, K., Frank, J., and Schulten, K. (2008) Flexible fitting of atomic structures into electron microscopy maps using molecular dynamics. Structure. 16, 673–683

48. Liebschner, D., Afonine, P. V., Baker, M. L., Bunkóczi, G., Chen, V. B., Croll, T. I., Hintze, B., Hung, L. W., Jain, S., McCoy, A. J., Moriarty, N. W., Oeffner, R. D., Poon, B. K., Prisant, M. G., Read, R. J., Richardson, J. S., Richardson, D. C., Sammito, M. D., Sobolev, O. V., Stockwell, D. H., Terwilliger, T. C., Urzhumtsev, A. G., Videau, L. L., Williams, C. J., and Adams, P. D. (2019) Macromolecular structure determination using X-rays, neutrons and electrons: recent developments in Phenix. Acta Crystallogr D Struct Biol. 75, 861–877

49. Emsley, P., Lohkamp, B., Scott, W. G., and Cowtan, K. (2010) Features and development of Coot. Acta Crystallogr. D Biol. Crystallogr. 66, 486–501

50. Chen, V. B., Arendall, W. B., 3rd, Headd, J. J., Keedy, D. A., Immormino, R. M., Kapral, G. J., Murray, L. W., Richardson, J. S., and Richardson, D. C. (2010) MolProbity: all-atom structure validation for macromolecular crystallography. Acta Crystallogr. D Biol. Crystallogr. 66, 12–21

51. Laskowski, R. A., Jabłońska, J., Pravda, L., Vařeková, R. S., and Thornton, J. M. (2018) PDBsum: Structural summaries of PDB entries. Protein Sci. 27, 129–134

52. Rigsby, R. E., and Parker, A. B. (2016) Using the PyMOL application to reinforce visual understanding of protein structure. Biochem. Mol. Biol. Educ. 44, 433–437

53. Pettersen, E. F., Goddard, T. D., Huang, C. C., Couch, G. S., Greenblatt, D. M., Meng, E. C., and Ferrin, T. E. (2004) UCSF Chimera--a visualization system for exploratory research and analysis. J. Comput. Chem. 25, 1605–1612

54. Hsu, S.-T. D., Fucini, P., Cabrita, L. D., Launay, H., Dobson, C. M., and Christodoulou, J. (2007) Structure and dynamics of a ribosome-bound nascent chain by NMR spectroscopy. Proc. Natl. Acad. Sci. U. S. A. 104, 16516–16521

