## Supporting Information for "Molecular basis for the ATPase-powered substrate translocation by the Lon AAA+ protease"

This file includes:

- Figs. S1 to S6
- Legend for Video S1
- Tables S1

Other Supporting Information for this manuscript includes the following:

- Video S1

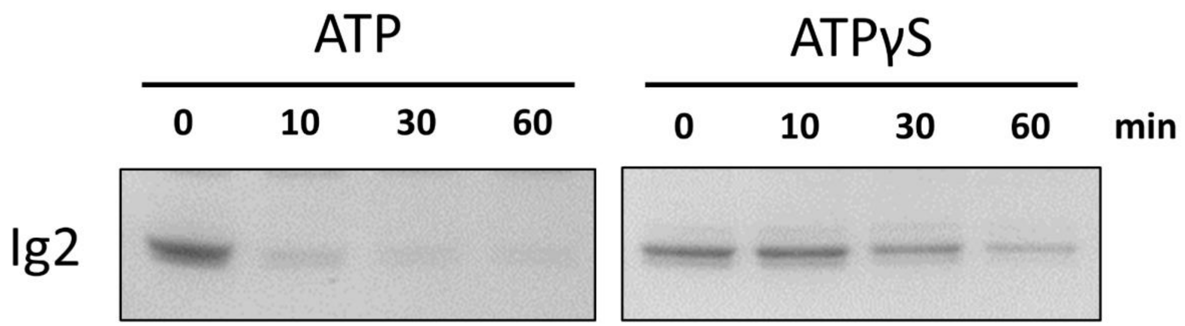


**Supporting Figure S1.** Substrate degradation assay. The substrate protein (Ig2) was degraded by wild-type MtaLonA in the presence of ATP (left) or ATP-γ-S (right).

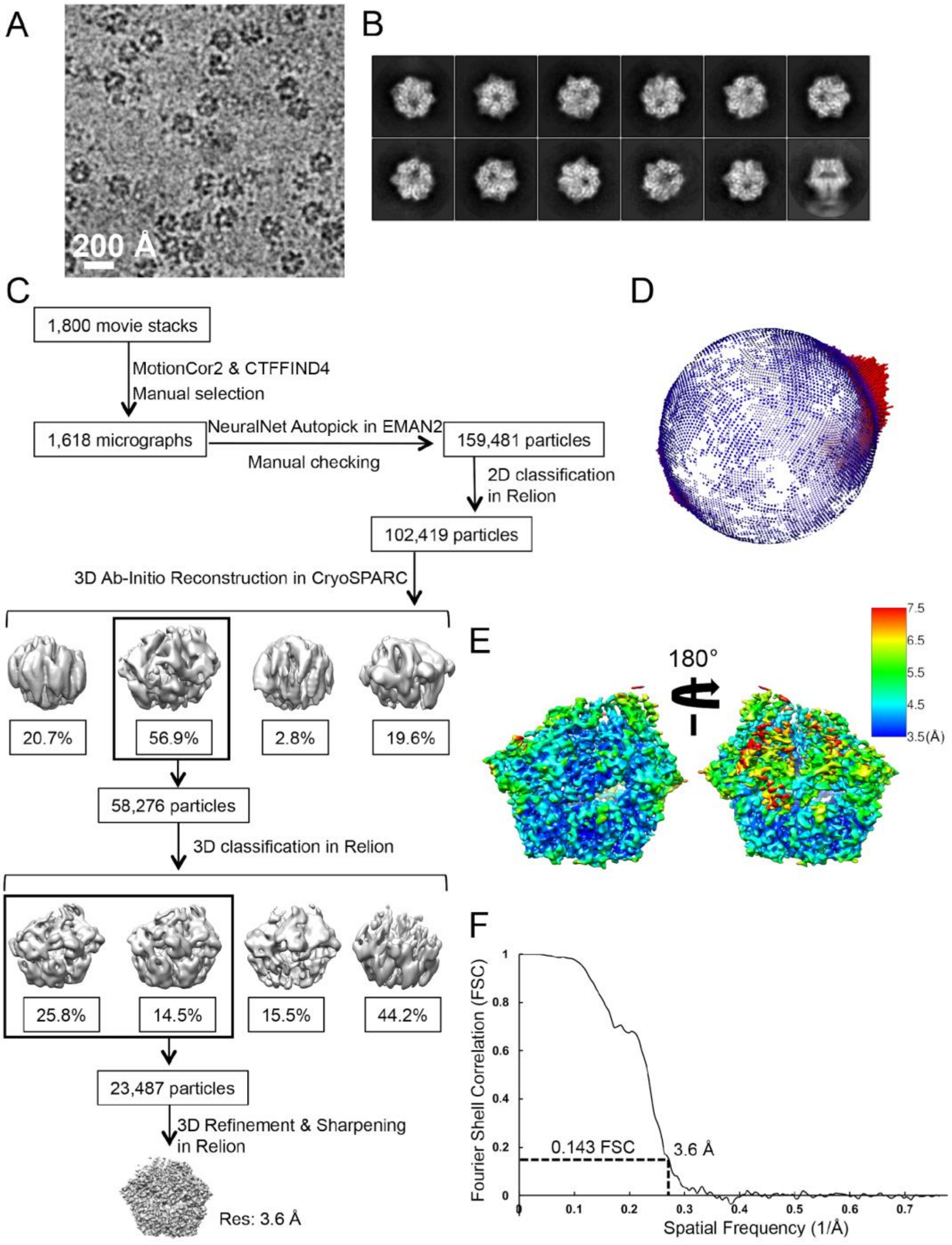


**Supporting Figure S2.** Single-particle cryo-EM analysis of the substrate-engaged MtaLonA. (**A**) Representative motion-corrected cryo-EM micrograph. (**B**) Reference-free 2D class averages. (**C**) Workflow of cryo-EM data processing. (**D**) Euler angle distribution of all particles used for calculating the final 3D reconstruction. (**E**) Resolution map for the final 3D reconstruction. (**F**) Gold standard FSC plot for the final 3D reconstruction.

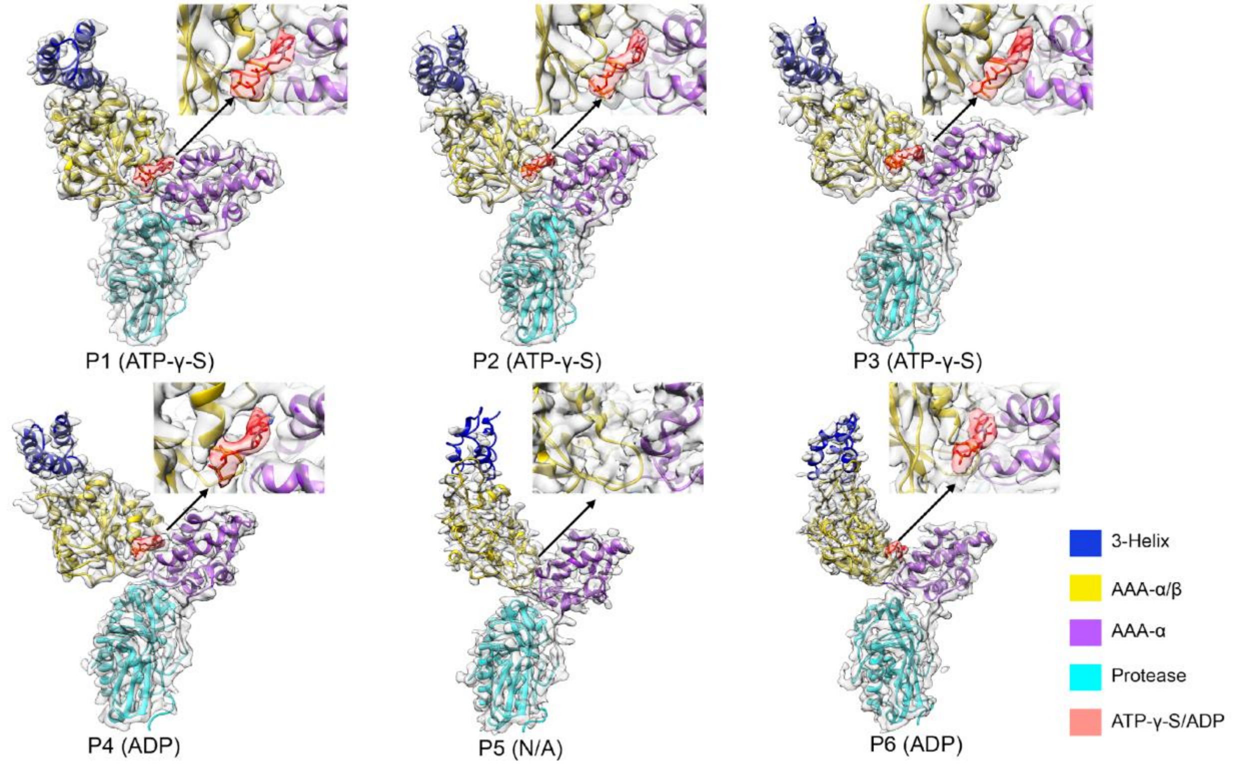


**Supporting Figure S3.** Identification of the bound nucleotide in the ATPase sites of protomers P1-P6. The nucleotide density with fitted model is shown in the insets. Different domains are shown in different colors.

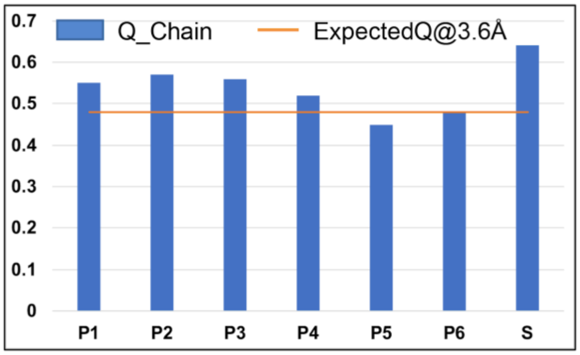


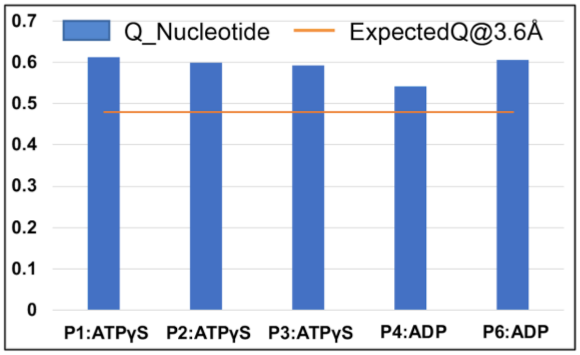


**Supporting Figure S4.** Resolvability of the cryo-EM structure of MtaLonA complexes calculated by Q-scores (29). The Q-scores are plotted for each chain and nucleotide in the model and the map; the orange line represents the expected Q-score at the respective resolution based on the correlation between Q-scores and map resolution of proteins. The higher Q-score indicates better resolvability. S refers to the substrate Ig2.

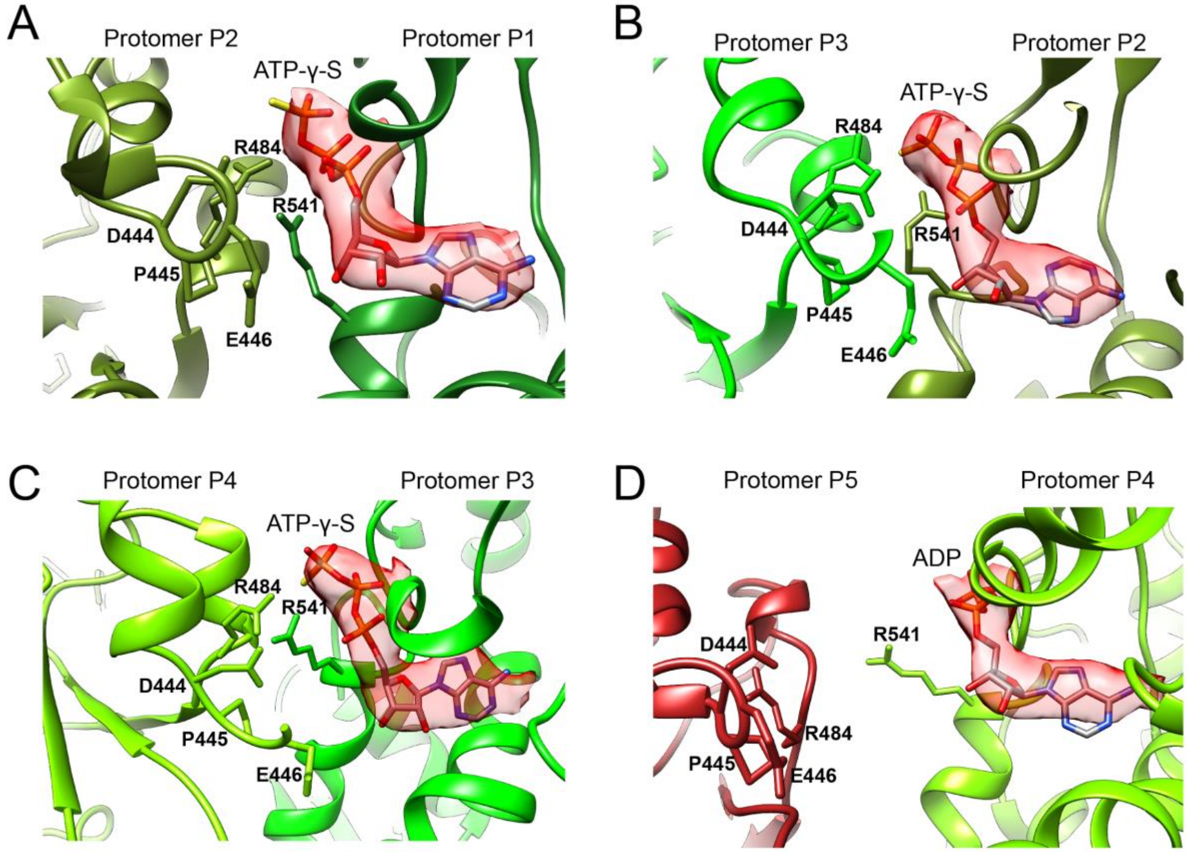


**Supporting Figure S5.** Conformation of the PS1βH base-loop (Asp444, Pro445, Glu446) and Arg-finger (Arg484) in the nucleotide-binding pocket between the neighboring protomers
The bound nucleotide is shown in red density with the model fitted in.

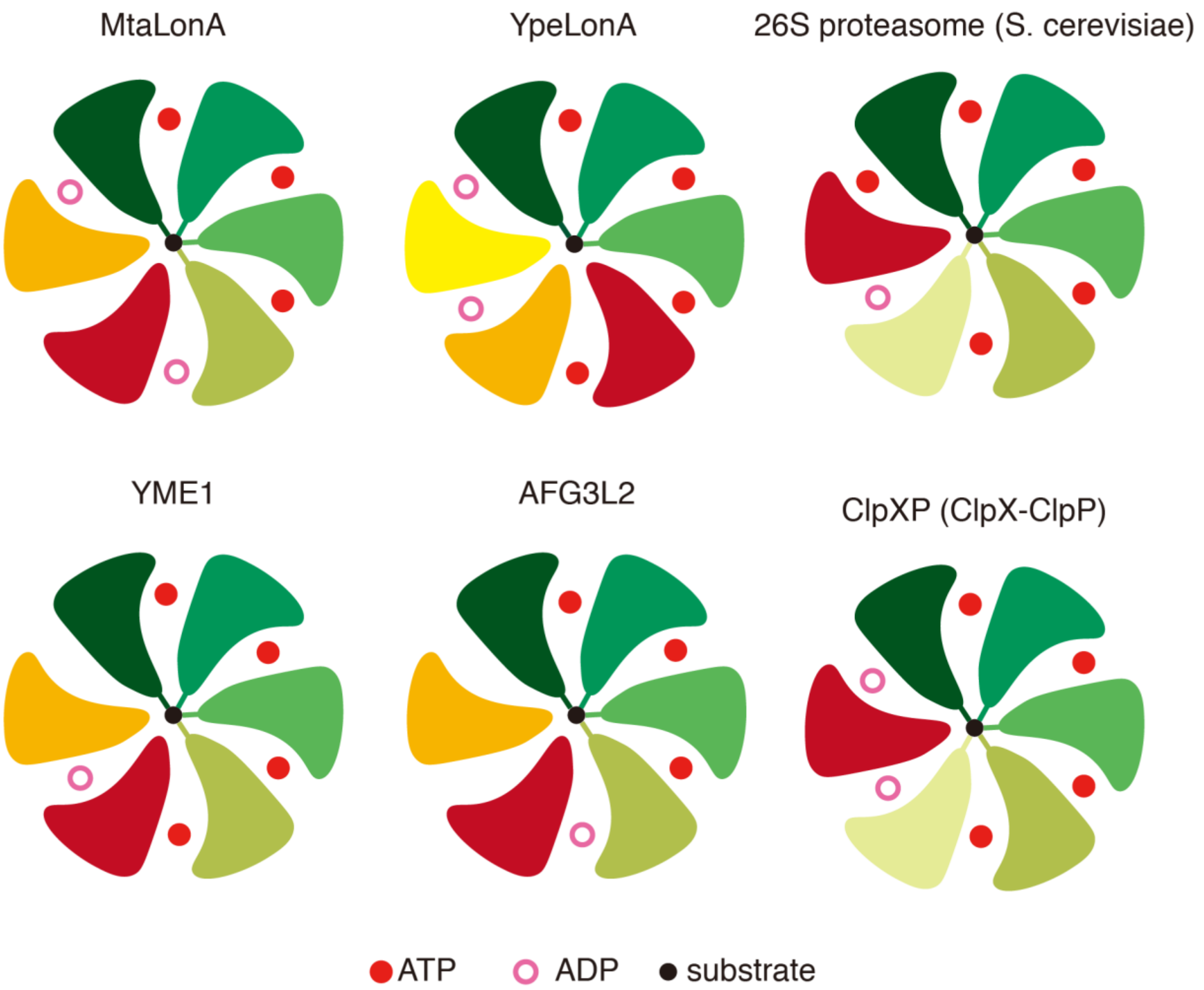


**Supporting Figure S6.** Comparison of the nucleotide-binding states for different AAA+ proteases bound to substrates. The top-view diagrams are shown to illustrate the binding states of the nucleotides and the substrate. Substrate-engaged protomers are colored in different shades of green.

**Supporting Video S1.** The cryo-EM map of the substrate-bound MtaLonA complex with the model fitted in. Four Tyr397 residues from the pore-loop 1 of protomers P1-P4 contact the substrate (Ig2) in a right-handed spiral arrangement. The nucleotides in the ATPase domain in protomers P6, P1, P2, P3, and P4 are shown in an amplified view sequentially.

**Supporting Table S1. Cryo-EM data collection, processing, and model validation**

| | Substrate-engaged MtaLonA |
|---|---|
| Data collection and processing | |
| Microscope | Titan Krios |
| Voltage (kV) | 300 |
| Camera | Gatan K2 Summit |
| Pixel size (Å) | 0.65 |
| Total Dose (e-/Å2) | 51 |
| Exposure time (s) | 6 |
| Number of frames per exposure | 30 |
| Defocus range (μm) | -0.6 - -2.6 |
| Number of micrographs | 1,800 |
| Number of initial particles | 159,481 |
| Number of particles for 3D analyses | 102,419 |
| Symmetry | C1 |
| Number of final particles | 23,487 |
| Resolution (0.143 FSC, Å) | 3.6 |
| Atomic model refinement | |
| Software | phenix.real_space_refine |
| Clash score | 21.78 |
| MolProbity score | 2.32 |
| Poor rotamers | 0.51% |
| Favored rotamers | 92.35% |
| Ramachandran outliers | 0.22% |
| Ramachandran favored | 91.81% |
| Bad bonds | 0.03% |
| Bad angles | 0.04% |
